# Niclosamide loaded Eudragit EPO nanoparticles show enhanced *Candida* biofilm penetration, trigger biofilm detachment and protect from mucosal candidiasis

**DOI:** 10.1101/2022.06.02.494588

**Authors:** Yogesh Sutar, Sunna Nabeela, Shakti Singh, Abdullah Alqarihi, Norma Solis, Teklegiorgis Ghebremariam, Scott Filler, Ashraf S. Ibrahim, Abhijit Date, Priya Uppuluri

## Abstract

*Candida albicans* biofilms are a complex multilayer community of cells that are resistant to almost all classes of antifungal drugs. The bottommost layers of biofilms experience nutrient limitation where *C. albicans* cells are required to respire. We previously reported that a protein Ndu1 is essential for *Candida* mitochondrial respiration; loss of *NDU1* causes inability of *C. albicans* to grow on alternative carbon sources and triggers early biofilm detachment. Here, we screened a repurposed library of FDA approved small molecule inhibitors, to identify those that prevent NDU1-associated functions. We identified an anti-helminthic drug, Niclosamide (NCL), which not only prevented growth on acetate, *C. albicans* hyphenation and early biofilm growth, but also completely disengaged fully grown biofilms of drug resistant *C. albicans* and *C. auris* from their growth surface. To overcome the sub-optimal solubility and permeability of NCL that is well-known to affect its *in vivo* efficacy, we developed NCL encapsulated Eudragit EPO (an FDA-approved polymer) nanoparticles (NCL-EPO-NPs) with high niclosamide loading, that also provided long-term stability. The developed NCL-EPO-NPs completely penetrated mature biofilms and attained anti-biofilm activity at low microgram concentrations. NCL-EPO-NPs induced ROS activity in *C. albicans*, and drastically reduced oxygen consumption rate in the fungus, similar to that seen in an NDU1 mutant. NCL-EPO-NPs also significantly abrogated mucocutaneous candidiasis by fluconazole resistant strains of *C. albicans*, in mice models of oropharyngeal and vulvovaginal candidiasis. To our knowledge, this is the first study that targets biofilm detachment as a target to get rid of drug-resistant *Candida* biofilms, and uses nanoparticles of an FDA approved non-toxic drug to improve biofilm penetrability and microbial killing.

## Introduction

*C. albicans* is a part of the normal human microbiota. In individuals with compromised immunity, the fungus typically infects target organs by overgrowth or hematogenous spread from colonized sites within the body. The incidence of hematogenously disseminated candidiasis in the United States is ~ 60,000 cases per year, making *Candida* spp. one of the most-frequently isolated nosocomial bloodstream pathogen^1–4^, carrying unacceptably high crude and attributable mortality rates of about 40-60%, despite antifungal drug treatment^5,6^. Of concern is the emergence of intrinsically drug resistant species of *Candida*, such as *C. auris* that can be refractory to all classes of FDA-approved antifungal drugs^7^.

The success of *Candida* spp. as a pathogen is predominantly due to the organism’s ability to adhere robustly to abiotic surfaces, filament and produce biofilm infections on medical devices^8^. A biofilm is a community of microbes attached to a surface and encased in an extracellular matrix^9–11^. Due to the sheer high density of cells^12^, and the ability of these cells to produce a thick extracellular matrix^11,13^, cells are shielded from environmental assaults, innate immune cells and from killing by antifungal drugs. Studies of catheter-related *Candida* infection have unequivocally shown that retention of vascular catheters is linked to prolonged fungemia, failure of antifungal therapy, increased risk of metastatic complications, and death (with mortality rates of >40%)^14–16^. Unfortunately, in many cases removal of catheters or implanted devices is not possible, therefore alternative strategies to manage biofilms are needed.

Such recalcitrance of biofilm cells is not unique to fungi, but also resonates throughout the bacterial biofilm setting. Antibiotics have often proven impotent against biofilms, and to date no antibiotic has been developed for use against biofilm infections^17^. The current most-promising focus to combat bacterial biofilms has been to unhinge and disperse them into a population of planktonic cells that would immediately lose properties of antibiotic resistance^17–20^. Such an approach may not be easy when it comes to *Candida* biofilms considering fungal biofilms are made up of a scaffold of hyphal elements that cannot be broken easily. However, an approach that could induce early detachment of biofilms might hold promise. Indeed, biofilm dispersed cells are more susceptible to antifungal drugs than parent biofilms^21^.

Biofilms experience nutrient limitations specifically within the inner layers of the community^22^. We recently reported that a conserved mitochondrial protein NDU1 is essential for *C. albicans* respiration and survival in non-fermentative conditions^23^. Interestingly, *C. albicans* NDU1 mutant strains could develop biofilms but these biofilms detach early from their growth substrate^23^. It was determined that the glucose-starved environment within the biofilm lead to increased ROS accumulation and low oxygen consumption rates in NDU1 mutant, culminating to the detachment phenotype. Here, we used NDU1 as a target to screen a repurposed library of FDA-approved small molecule inhibitors (New Prestwick Chemical library) to isolate those that could prevent *C. albicans* growth in acetate. We identified Niclosamide (NCL), which has a long history of safety and efficacy as an anthelmintic drug in children and adults. When used against *Candida*, NCL treatment was found to phenocopy the defects of an NDU1 mutant.

NCL is a weakly acidic (pK_a_: 6.87) hydrophobic drug with very low solubility in water and gastrointestinal fluids, displays poor permeability and intestinal glucuronidation, which sorely limit its oral bioavailability and *in vivo* efficacy ^24^. To overcome this drawback, we encapsulated NCL in nanoparticles (NPs) of an FDA-approved cationic polymethylmethacrylate polymer called Eudragit EPO (EPO), to enhance its bioavailability and *in vivo* efficacy. Here, we elucidate that NCL can stably be encapsulated into EPO NPs with high loading due to the molecular-level interaction between cationic EPO and weakly acidic NCL. NCL loaded EPO NPs (NCL-EPO-NPs) show long-term physicochemical and chemical stability, inhibit growth and filamentation of *C. albicans* as well as *C. auris*, prevent biofilm growth (at concentrations as low as 0.5-2 µg/ml), and most importantly, disengage preformed biofilms from their growth material. Furthermore, incorporation of NCL-EPO-NPs into an in-situ thermogelling formulation facilitates their intra-oral delivery and shows significantly potent antifungal activity in two independent mouse models of mucosal infections.

This is the first study targeting the process of biofilm detachment for identification of an anti-biofilm inhibitor. Furthermore, successful repurposing of an FDA-approved inhibitor in a nanoparticulate form for treatment against *C. albicans* has not been described before. In summary, we have targeted a novel *Candida* protein (NDU1) using nanoformulation of an FDA-approved antiparasitic drug, facilitating its rapid repurposing and development as an urgently needed antifungal agent.

## Results

### NCL targets mitochondrial respiration and induces biofilm detachment

Using a forward genetic screening approach, we recently identified and reported on *NDU1*, a gene controlling *C. albicans* biofilm dispersal and biofilm detachment^23^. NDU1 is a mitochondrial inner membrane protein required for respiration via Complex I of the electron transport chain, is essential for growth on alternative carbon sources, and required for virulence *in vivo*^23^. Our work highlighted the connection between respiration and biofilm sustenance, where impaired mitochondrial respiratory activity in an *NDU1* mutant leads to reduction in biofilm dispersal and early biofilm detachment. As has been well-documented in the bacterial biofilm field^17–20^, we postulated that targeting early biofilm detachment is a promising approach to combat biofilm-associated infections. To test this hypothesis, we used NDU1 as a target to identify small molecule inhibitors that would inhibit NDU1-associated functions and consequently trigger biofilm detachment.

*C. albicans* lacking *NDU1* are inviable in media containing acetate (or any other non-fermentative carbon sources), but grow as robustly as the wild type (WT) in media with glucose^23^. We harnessed this unmistakable phenotype to screen a library of 1200 repurposed FDA-approved small molecule inhibitors against a *C. albicans* strain overexpressing NDU1 (OE) in media containing 1% acetate. An inhibitor that could at a low concentration of 10 µM, curtail biofilm growth of OE in acetate, but not of OE or the WT in glucose was considered a “hit”. This ensured the recognition of an inhibitor that only targets NDU1 and not general viability per se.

A total of 21 inhibitors were identified that prevented >50% of OE in acetate (hit rate of 1.7%; see S1). However, 15 out of the 21 were either disinfectants (e.g. hexachlorophene, thimerosal) or known antifungal drugs (e.g. fluconazole, tioconazole, econazole etc.) that were inhibitory to *C. albicans* growth even in glucose containing medium (S1) and hence were discarded. We identified antimycin A, a known inhibitor of oxidative phosphorylation as a hit in acetate and not glucose, corroborating our assay that focused on identifying inhibitors of cellular respiration. However, antimycin A is considered extremely toxic and hence disregarded as a molecule of interest. Additional hits were alexidine dihydrochloride and auranofin which we^25^ and others^26^ have previously shown to be attractive molecules for antifungal development, and are currently under different stages of drug development. Nevertheless, these drugs inhibited general viability rather than respiration and were rejected for this study. Finally, we identified NCL, an anti-helminthic drug, as the only inhibitor that curtailed biofilm growth of NDU1 OE strain in acetate and not glucose, thereby potentially targeting NDU1 activity, or affecting respiratory chain in a similar manner.

### Eudragit EPO (EPO) and NCL show concentration-dependent molecular interaction leading to the stabilization of NCL in the EPO matrix

NCL suffers from major physiochemical drawbacks such as high hydrophobicity, poor permeability and intestinal glucuronidation which leads to poor bioavailability, thereby greatly limiting its widespread clinical applications. Nanoparticles based drug delivery systems have the potential to solubilize hydrophobic drugs like NCL, thus improving permeability and *in vivo* applicability of the drug^27^. To identify the nanoformulation polymers suitable for the development of NCL nanoparticles, we first evaluated pharmaceutically accepted anionic (polyvinyl acetate phthalate and Eudragit S100) and cationic (EPO) polymers for their ability to stabilize NCL in the polymer matrix and to facilitate the development of NCL nanoparticles. Our simple solvent-antisolvent precipitation method involving a fixed ratio of NCL and polymers showed that only the EPO-NCL combination could form a stable dispersion. To ascertain the stabilization effect of EPO seen in our screening studies, we evaluated solid-state interaction between NCL and EPO at various ratios (1:10, 2:10, 3:10, and 4:10 w/w) using Fourier-transform infrared (FT-IR) spectroscopy and nuclear magnetic resonance (NMR) spectroscopy. Our FT-IR spectroscopic studies showed the characteristic peak for –OH and/or N-H (amide) stretching of NCL at 3235 cm^−1^, while O-H bending and C-N stretching vibrations appeared at ~1340 and 1280 cm^−1^ respectively (**Fig.1A,B**). Interestingly, FT-IR spectra of NCL-EPO coevaporates showed disappearance of −OH and/or N-H stretching vibrations of NCL irrespective of the weight ratio (**Fig. 1A**) whereas O-H bending and C-N stretching vibrations of NCL showed concentration-dependent red shift (**Fig. 1B**). Additionally, peaks corresponding to dimethylamino group^28^ (2770 & 2820 cm^−1^) and the carboxyl group of ester (1723 cm^−1^) in EPO showed concentration-dependent reduction in the peak intensity along with the red shift in NCL-EPO coevaporates (**Fig.1A,B**). The changes in FTIR spectra of NCL-EPO coevaporates compared to individual components indicate the strong concentration dependent molecular interaction between NCL and EPO. The spectral changes suggest that hydrogen donating groups in NCL (O-H & N-H) are stabilized by hydrogen accepting groups in EPO (C=O, dimethylamino)^29,30^ in a concentration-dependent manner^31–33^.

**FIGURE 1.**
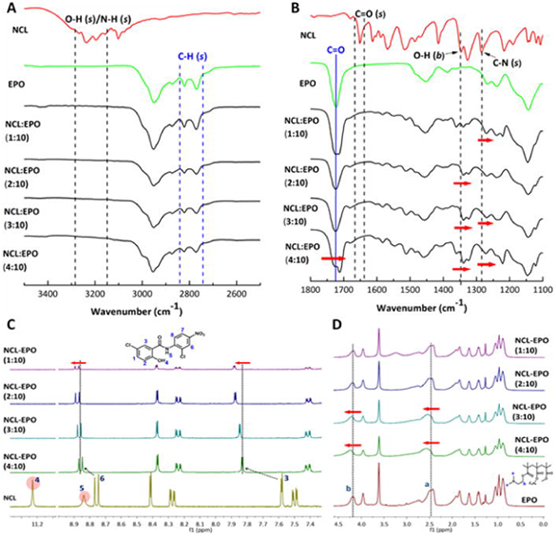
Niclosamide (NCL) interacts with Eudragit EPO (EPO) at a molecular level and the concentration of NCL relative to EPO impacts the extent of molecular interaction. Solid dispersion containing different weight ratios of NCL and EPO (1:10, 2:10, 3: 10 and 4:10) were prepared using the solvent evaporation method. Fourier-transform infrared (FT-IR) spectroscopic studies (A) and (B) showed that −OH and/or NH stretching vibrations of NCL abolished in all the NCL-EPO solid dispersions irrespective of the weight ratio indicating strong interaction between NCL and EPO. Furthermore, −OH bending vibrations, and C-N stretching vibrations of NCL, C=O stretching vibrations and −CH stretching vibrations of EPO showed concentration dependent redshift and diminution in the peak intensity indicating the effect of NCL to EPO weight ratio. The nuclear magnetic resonance (NMR) spectroscopy studies (C) and (D) showed that proton corresponding to −OH (marked with number 4) and −NH (marked with number 5) in NCL abolished irrespective of the NCL to EPO weight ratio indicating a strong electrostatic interaction. Additionally, signals for the aromatic protons from NCL and aliphatic protons from EPO showed concentration dependent downfield shift further confirming the interaction between NCL and EPO. The spectroscopic studies indicated the possibility of achieving high loading and stabilization of NCL in the EPO matrix.

We further validated the molecular interaction between NCL and EPO using NMR spectroscopy. The NMR spectra of all NCL-EPO coevaporates showed disappearance of the protons corresponding to O-H and N-H functional group in NCL irrespective of the NCL-EPO ratio whereas the protons corresponding to the tertiary amine of EPO (dimethylamino) showed a downfield shift in NCL:EPO ratios 3:10 and 4:10 and decrease in the signal peak intensity at all NCL:EPO ratios, indicating strong hydrogen bond formation between O-H/N-H group of NCL and tertiary amine of EPO ^31^ (**Fig. 1C & D**). Additionally, NMR spectra of NCL-EPO coevaporates showed a concentration-dependent downfield shift in the signal of aromatic protons of NCL. It is likely that the cationic center and long polymeric chain of the EPO could influence the electron density of the adjacent methylene units in NCL causing the proton shifts ^34^. Thus, NMR spectroscopic studies corroborated the inferences from FT-IR studies. Taken together, spectroscopic studies confirmed the ability of EPO to stabilize NCL via molecular level interaction indicating a possibility of developing EPO NPs with a potentially high concentration of NCL.

### Eudragit EPO allows for stable encapsulation, high loading, and long-term stability of NCL in nanoparticles

We evaluated various FDA-approved surfactants in a simple and scalable nanoprecipitation method for the development of NCL loaded EPO NPs (NCL-EPO-NPs). Surfactants such as Kolliphor RH 40 (RH 40), Poloxamer 407 (P407), Vitamin E TPGS (TPGS), Kolliphor ELP (ELP), Polysorbate 20 (PS 20): Polysorbate 80 (PS80), Kolliphor HS 15 (HS 15) were evaluated for their ability to yield NCL-EPO-NPs with lowest particle size, acceptable polydispersity index and colloidal stability of at least 1 day (for initial screening). The size, polydispersity, and colloidal stability of NPs varied depending upon the type of surfactant used (**Fig 2A**). NCL-EPO-NPs stabilized by P407 showed optimal particle size, polydispersity, and colloidal stability and were selected for further development.

**FIGURE 2.**
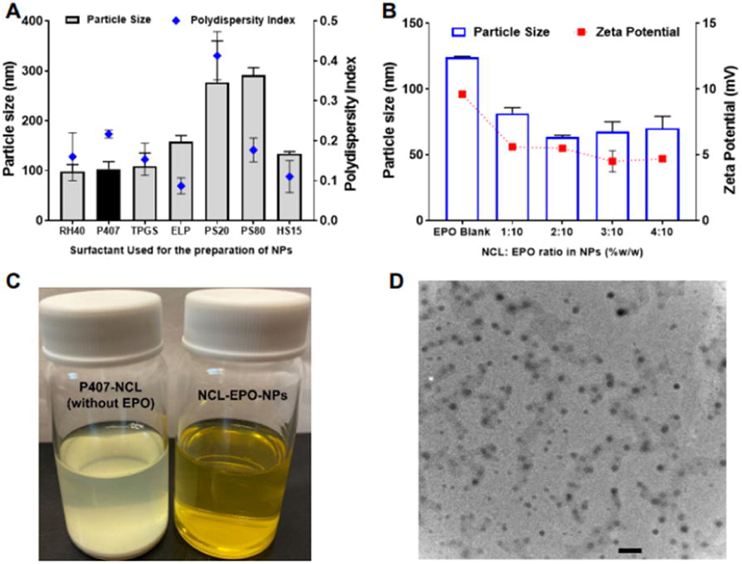
Development of Poloxamer 407 stabilized Eudragit EPO nanoparticles with high niclosamide loading (NCL-EPO-NPs). (A) Screening of various FDA-approved surfactants for the development of NCL loaded EPO NPs. RH40: Kolliphor RH 40; P407: Poloxamer 407; TPGS: Vitamin E TPGS; ELP: Kolliphor ELP; PS 20: Polysorbate 20; PS80: Polysorbate 80; HS 15: Kolliphor HS 15. P407 stabilized NCL-EPO-NPs were selected for the further characterization and studies. Data expressed as mean ± S.D. (n =3); (B) Effect of NCL loading on the size and zeta potential of EPO NPs. It was possible to obtain stable NCL-EPO-NPs with high NCL loading of 40% w/w of EPO. The polydispersity index value for all samples was < 0.26. Data expressed as mean ± S.D. (n =3) (C) A representative image to demonstrate critical role of Eudragit EPO for the formation of stable NCL-EPO-NPs. In the absence of Eudragit EPO, niclosamide immediately precipitated indicating the importance of Eudragit EPO. (D) Transmission electron microscopy (TEM) image of P407-NCL-EPO-NPs. The NPs showed spherical morphology and a reasonable correlation with the dynamic light scattering data. (Scale bar = 200 nm)

Thus, we evaluated the feasibility of developing P407 stabilized EPO-NCL-NPs with higher loading of NCL. It was possible to develop P407 stabilized EPO-NCL-NPs at NCL:EPO ratios of 1:10, 2:10, 3:10, and 4:10 (**Fig 2B**) indicating the feasibility of developing NCL-EPO-NPs at high NCL loading.

Compared to blank EPO NPs, NCL-EPO-NPs showed lower particle size and surface charge which further confirmed the molecular interaction between NCL and EPO in the nanoparticulate form. The UV spectrophotometric analysis (**S2 A,B**) showed that the encapsulation efficiency of NCL was > 98% in NCL-EPO-NPs irrespective of the NCL:EPO ratio used for the nanoformulation. NCL-EPO-NPs with NCL loading of 40% (%w/w compared to EPO) showed particle size of 70.2 ± 9 nm, the average polydispersity of 0.23, the average zeta potential of +4.5 mV, and excellent colloidal stability (**Fig 2 A,B**). Any attempts to prepare NCL NPs without EPO immediately led to the precipitation of NCL further confirming the critical role of EPO in the stable encapsulation of NCL (**Fig 2C**). The TEM imaging showed that EPO-NCL-NPs have spherical morphology (**Fig 2D**) and their size was in congruence with the size determined by the dynamic light scattering.

We carried out several studies to confirm the molecular interaction between NCL and EPO at the nanoscale. First, we evaluated the fluorescence spectra of NCL solution and NCL-EPO-NPs aqueous dispersion using a spectrofluorometer. The hydroalcoholic NCL solution showed maximum emission at 406 nm, whereas NCL-EPO-NPs dispersed in water showed maximum emission at 526 nm (**Fig 3A**). The significant red shift and no increase in the fluorescence quantum yield observed in the case of NCL-EPO-NPs indicate strong NCL-EPO interaction and stable encapsulation of NCL at the nanoscale. Further, we freeze dried NCL-EPO-NPs and evaluated the molecular interaction of NCL and EPO in NPs using FT-IR and NMR spectroscopy. As anticipated, the FTIR spectroscopic analysis of freeze dried NCL-EPO-NPs showed the absence of –OH and –NH stretching vibrations of NCL and shifts in the C=O stretching vibration of EPO indicating the retention of molecular interaction between NCL and EPO in the nanoparticles (**Fig 3B**). Interestingly, a shift in C-O stretch (ether) of P407 (1101 cm^−1^) was also observed in the freeze dried NCL-EPO-NPs-FD (1090 cm^−1^) suggesting the molecular interaction between P407 and NCL-EPO which may further help in the stabilization of NCL-EPO NPs (**S2C**). The changes in the NMR spectrum of freeze-dried NCL-EPO-NPs compared to pure NCL and EPO were similar to that observed in the NCL-EPO coevaporates (described previously), which further confirmed the retention of strong molecular interaction between NCL and EPO in the nanoparticles (**Fig 3C; S2D**).

**FIGURE 3.**
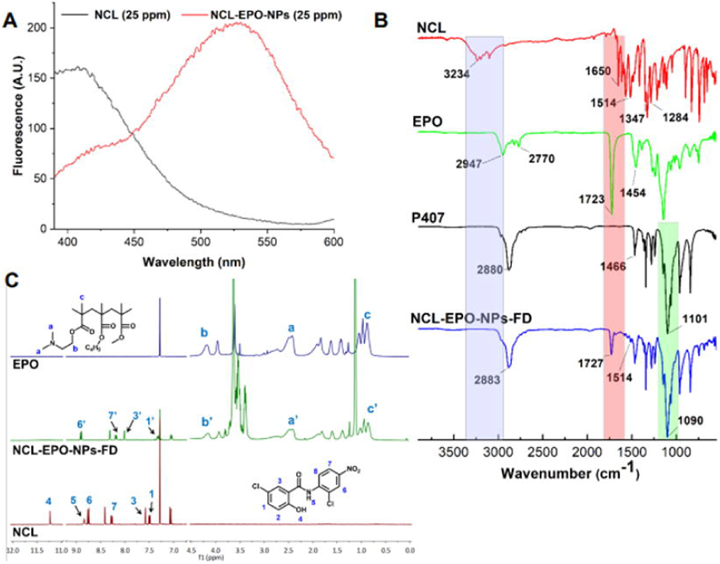
Niclosamide-Eudragit EPO molecular interaction is maintained in the nanoformulation. P407 stabilized NCL-EPO-NPs with NCL loading of 40% w/w of EPO were either used as an aqueous dispersion or freeze dried to obtain dry powder (NCL-EPO-NPs-FD) for FT-IR and NMR analysis. (A) Spectrofluorometric analysis of NCL hydroalcoholic solution (25 ppm) and aqueous dispersion of NCL-EPO-NPs (equivalent to 25 ppm NCL) showed a significant red shift in the case of NCL-EPO-NPs indicating strong molecular interaction between NCL and EPO in the nanoparticles. (B) FT-IR spectroscopic analysis of NCL-EPO-NPs-FD showed absence of −OH and −NH stretching vibrations of NCL and shifts in the C=O stretching vibrations of EPO indicating the retention of molecular interaction between NCL and EPO in the nanoparticles. (C) NMR spectroscopic analysis of NCL-EPO-NPs-FD showed disappearance of protons corresponding to −OH and −NH functional groups in NCL indicating retention of strong molecular interaction between NCL and EPO in the nanoparticles.

It is imperative that the nanoformulation developed after several optimization and characterization experiments maintain long-term physical (size, polydispersity index, surface charge, and pH) and chemical (NCL content) stability. The optimized P407 stabilized NCL-EPO-NPs with different NCL loading (% w/w of EPO) were evaluated for physical and chemical stability for a period of 4 weeks. NCL-EPO-NPs with NCL loading of 40% (% w/w of EPO) showed negligible changes in the particle size, polydispersity, surface charge, and pH over a period of 4 weeks (**Fig 4 A,B**). The fluorescence spectra of NCL from NCL-EPO-NPs showed no to negligible changes even after 4 weeks (**Fig 4C**) indicating stable encapsulation and chemical stability of NCL in the nanoparticles. The 4-week stability data for NCL-EPO-NPs with lower NCL loading, and other characteristics are included in the supplementary section (**S2E**). Thus, we successfully developed P407 stabilized NCL-EPO-NPs with high loading of NCL that have long-term physical and chemical stability.

**FIGURE 4.**
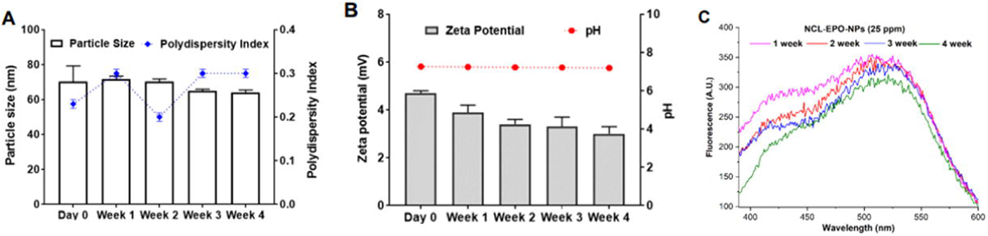
P407 stabilized NCL-EPO-NPs with NCL loading of 40% w/w of EPO show physical and chemical stability for 4 weeks. Three different batches of NCL-EPO-NPs were prepared, and their physical and chemical stability was evaluated for 4 weeks. (A) Particle size and polydispersity index of NCL-EPO-NPs over a period of 4 weeks showed negligible changes. Data expressed as mean ± S.D. (n =3); (B) Zeta potential and pH of NCL-EPO-NPs did not show any appreciable changes after 4 weeks of storage. Data expressed as mean ± S.D. (n =3); (C) The fluorescence intensity of NCL-EPO-NPs (NCL concentration: 25 ppm) showed negligible changes for 4 weeks indicating chemical stability. Analysis was carried out in triplicate and representative fluorescence spectra are shown in the figure.

### NCL-EPO-NP display high efficacy against *C. albicans* and *C. auris* planktonic as well as biofilm growth

We first validated NCL-EPO-NP in a dose response assay against *C. albicans* planktonic yeast cells in 96 well plates, growing in media with glucose or with acetate. While NCL-EPO-NP prevented >80% growth of *C. albicans* in acetate at 1 µg/ml (3 µM), it was not effective against *C. albicans* viability in glucose (**Fig 5A**). Viability measured by spot testing (**Fig 5B)** and XTT assay (**Fig 5C**) provided a visual and a quantifiable measure of the outcome, respectively. These studies, indicated that NCL-EPO-NP was fungicidal at 0.5-1 µg/ml in acetate, however ineffective in glucose. The effect of the molecule on *C. albicans* growth in glucose was also pictured by microscopy which clearly revealed a gradual decrease in *C. albicans* proliferation over dose escalation in acetate, whereas a steady increase in growth in media with glucose (**S3A**). Importantly, we found that while the nanoparticles had no effect on *C. albicans* viability in glucose, it potently inhibited *C. albicans* filamentation at concentrations as low as 0.5 µg/ml (1.5 µM) (**Fig 5D**). Since NDU1 was used as a target to identify niclosamide, we questioned if *C. albicans* NDU1 heterozygous strain or its deletion mutant will be more susceptible to NCL-EPO-NP. Certainly, at a concentration as low as 1 µM, both *NDU1/ndu1* as well as *ndu1/ndu1* exhibited a 29.8% (±1.5%) and 34.5% (±1%) reduction in viability, respectively. This reduction in viability worsened to 63.7% (±4.4%) for the heterozygote and 75.7% (±1.4%) for the homozygote after treatment with 20 µg/ml of NCL-EPO-NP. Remarkably, the drug had no effect on wild type *NDU1/NDU1* (0% loss in viability) even at concentrations >80 µg/ml. The NCL-EPO-NP-induced haploinsuffciency of *NDU1* suggests that it is required to tolerate and is a target of Niclosamide.

**FIGURE 5.**
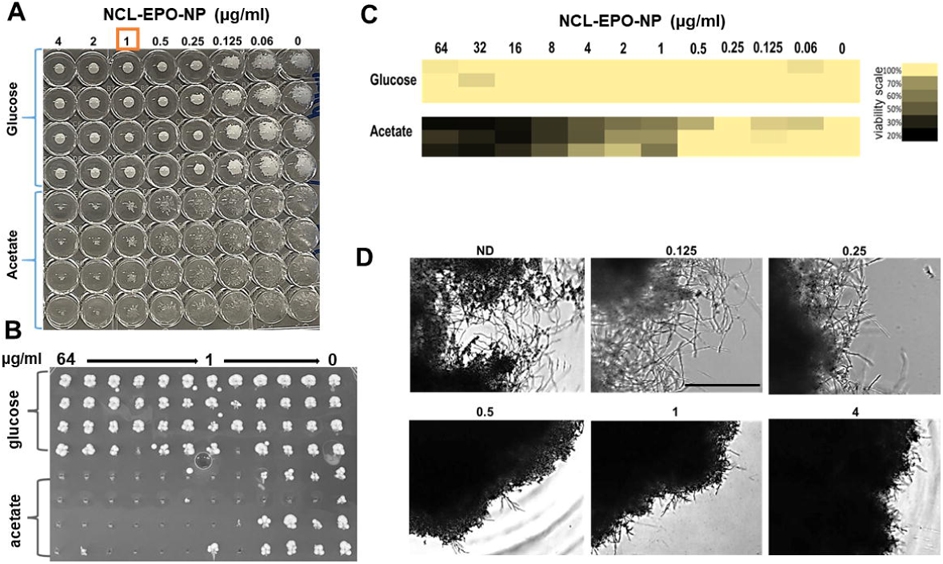
NCL-EPO-NP prevent filamentation and viability of planktonic *C. albicans* in glucose and acetate respectively. (A) Dose response assay using round bottom 96 well microtiter plates showed that NCL-EPO-NP could prevent growth of *C. albicans* in YP+1% acetate (MIC50 of 1 µg/ml). Also the NP appear to prevent filamentation of the fungus at 0.25 µg/ml in YP+1% glucose. (B) The viability was tested by spotting the NCL-EPO-NP treated cells on YPD agar. (C) Viability was also monitored using the XTT assay that measures the metabolic activity of cells. (D) Microscopic visualization of reduction in *C. albicans* filamentation on treatment with different concentrations of NCL-EPO-NP in the presence of 1% glucose. Scale bar = 500 µm.

Hyphal growth in *C. albicans* is vital for tissue invasion. Hence, we tested if the defect in filamentation caused by NCL-EPO-NP could block damage of human umbilical vascular endothelial cells (HUVEC) by *C. albicans*. Indeed, we found that the NCL-EPO-NPs could prevent 50% damage to HUVEC cells at doses between 2 and 4 µg/ml of NCL. NCL itself did not significantly damage the host cells even at 8 µg/ml (**S3B**).

Considering that filamentation is also a property imperative for biofilm growth, we investigated the effect of NCL-EPO-NP on *C. albicans* biofilm formation and on pre-formed biofilms. Again, NCL-EPO-NP prevented *C. albicans* biofilm growth when added at the time of biofilm initiation in RPMI - a pro-filamentation media that contains glucose. Biofilm growth was interrupted at 1 µg/ml, the dose that limits hyphal growth in the fungus (**Fig 6A**). In fact, when fully formed biofilms were treated for 24 h with NCL-EPO-NP, the biofilms were found to disintegrate and detach from the wells of the 96 well plate, leading to an overall decrease in biofilm biomass as measured by the XTT assay (**Fig 6B, C**). Such striking activity of a small molecule against preformed biofilms is uncommon, especially since it does not affect cell viability. Moreover, antifungal drugs have been reported to get sequestered by the biofilm ECM, which nullifies their potency. Thus, we tested the biofilm penetrability of the EPO-NPs stained with coumarin 6 which gave the NP a green color. We found that the nanoparticles could diffuse into the compact biofilm entity (hyphae stained red with Con A) all the way to the bottom of the biofilm cells (**Fig 6D, S3C**). Overall, our data demonstrate that NCL-EPO-NPs inhibit *C. albicans* hyphal growth, which has broader implications on virulence in terms of reduced invasion and biofilm growth, resulting in containment of infection and ridding of drug resistant biofilms.

**FIGURE 6.**
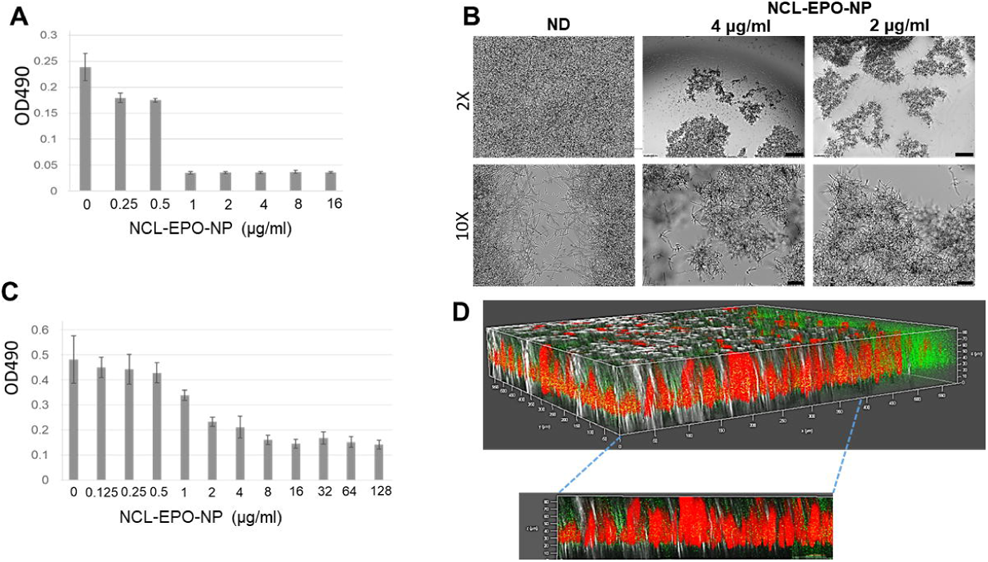
NCL-EPO-NP abrogate *C. albicans* biofilm growth. (A) Addition of NCL-EPO-NP at the beginning of C. albicans biofilm induction (96 well flat well microtiter plates), arrested biofilm growth at 1 µg/ml. (B) NCL-EPO-NP added on to preformed biofilms induced biofilm detachment at 2 µg/ml which was visualized microscopically. Scale bar for 2X mag is 500 µm, and for 10X is 100 µm. (C) The effect of the NP on preformed biofilms was quantified by XTT assay which revealed a >50% reduction in biofilm metabolic activity starting at 2 µg/ml. (D) Coumarin tagged NCL-EPO-NP (green) could efficiently penetrate the concanavalin stained *C. albicans* biofilm (red), as seen by the yellow stained particles (green+red overlap).

We next questioned if NCL-EPO-NP could also display similar efficacies against drug resistant *C. albicans* isolates or against the multidrug resistant fungus *C. auris*. Interestingly, we found that NCL-EPO-NP exhibited inhibitory activity exceeding 80% against two strains of *C. auris* in media containing glucose or acetate again between low concentrations of 0.5-1 µg/ml (**Fig 7A**). This meant the molecule was more potent against *C. auris* than *C. albicans* in glucose in which it only prevented filamentation. In fact, similar doses also inhibited growth of *C. auris* biofilms as measured by the XTT assay (**Fig 7B**) and when visualized by microscopy which showed inhibition of biofilm growth at 1 µg/ml (**Fig 7C**). Furthermore, the same concentration of NCL-EPO-NP was also efficacious at inhibiting >50% of biofilm growth of two most fluconazole resistant *C. albicans* strains (fluconazole MIC>128 µg/ml) and *C. auris* (fluconazole MIC>512) (**Fig 7D**).

**Figure 7:**
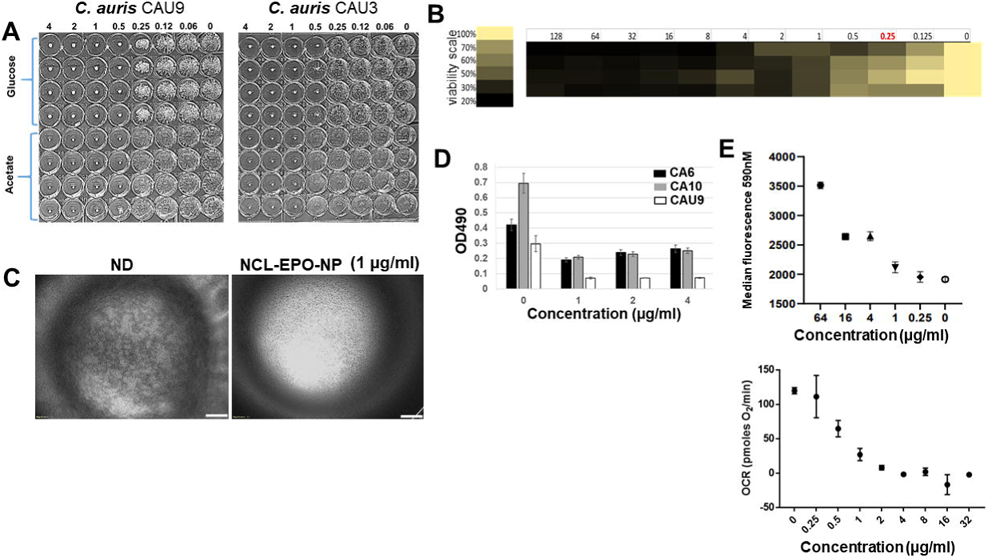
NCL-EPO-NP prevent growth of planktonic *C. auris*, increase ROS activity and negatively affect oxygen consumption rates in *C. albicans*. (A) Dose response assay using round bottom 96 well microtiter plates showed that NCL-EPO-NP could prevent growth of two different strains of *C. auris* (CAU-9 and CAU-3) in YP+1% acetate as well as in YP+1% glucose at concen-tration of 0.5-1 µg/ml. (B) The same dose 1 µg/ml and above, could also prevent growth of *C. auris* biofilms, measured by XTT reduction assay, and portrayed in a heat map where darker colors indi-cate increased fungal killing while yellow means fungal viability (4 replicates are shown). (C) NCL-EPO-NP at 1 µg/ml inhibited *C. auris* biofilm formation (bottom of the 96 well was microscopically visualized 2X mag. Scale bar = 500 µm). (D) NCL-EPO-NP at low concentration could prevent biofilm growth of flucon-azole resistant *C. albicans* strains CA-6, CA-10 and fluconazole resistant *C. auris* CAU-9. Results are from XTT assay with three replicates. (E) Increasing concentrations of NCL-EPO-NP correspondingly in-creased ROS production in *C. albicans*, which was measured by staining with MitoSox red with the resulting fluorescence read at 590nM by using a flow cytometer. (F) Increasing concentrations of NCL-EPO-NP also abrogated the oxygen consumption rate in *C. albicans*, which was measured using a Sea horse assay.

### NCL-EPO-NP increase intracellular ROS and reduces oxygen consumption rate in *C. albicans*

NCL was identified as the drug that potentially targets a mitochondrial protein NDU1 and negatively impacts an important aspect of cellular respiration such as the ability of *C. albicans* to grow on alternative carbon sources. We further tested its effect on other key features of mitochondrial respiration such as ROS production and mitochondrial oxygen consumption rates. Presence of NCL-EPO-NP significantly increased ROS accumulation in a dose dependent manner, with an 11% increase at 1 µg/ml, 32% increase at 4 µg/ml and 16 µg/ml and ~70% elevation at 64 µg/ml (**Fig 7E, S3D**). The fact that cells were hypersusceptible to NCL-EPO-NP in acetate indicated a faulty electron transport chain (ETC) activity ^23,35^. To test this hypothesis, we carried out Seahorse assays to test the respiratory prowess of *C. albicans* in DMEM medium without glucose, and with NCL. The cells presented a significant decrease in respiratory capacity with increasing doses of the NCL-EPO-NP. The oxygen consumption rate of cells fell below 50% between 0.5-1 µg/ml, and reduced to >90% by 2 µg/ml (**Fig 7F, S4A**), thus corroborating the findings of NCL MIC in media lacking glucose (**Fig 5A-C**). Together these results indicate that NCL-EPO-NP target mitochondrial respiration by interfering with the ETC.

### NCL-EPO-NP protects mice from mucosal infections

The pronounced *in vitro* efficacy of NCL-EPO-NP against *C. albicans* filamentation, invasion and biofilm growth encouraged us to test this molecule *in vivo*. To facilitate this, we developed a P407-based in-situ gelling formulation containing NCL-EPO-NPs. Following a strategy previously reported by us ^36,37^, NCL-EPO-NPs were incorporated in an in-situ gelling formulation composed of 20% w/v P407 and 1% w/v Poloxamer 188 (**S4B**). Incorporation of the NCL-EPO-NP in a gel form allowed us to facilitate their delivery directly into mucosal organs of mice.

We evaluated NCL-EPO-NP first in the established mouse model of oropharyngeal candidiasis (OPC) ^38,39^. Mice pretreated with corticosteroids were infected sublingually with *C. albicans*. After allowing the infection to establish for 2 d, cohorts of mice were treated with fluconazole or NCL-EPO-NP gel. Fluconazole was administered systemically, whereas the drug-containing gel was introduced into the mouth of the mice. Antifungal activity was assessed by measuring residual colony forming units (CFU) persisting on the tongue after completing 4 d of therapy. NCL-EPO-NP gel treatment decreased fungal burden by almost one log, compared to untreated mice and mice treated with placebo gel without NCL (**Fig 8A**). We further investigated the potency of NCL-EPO-NP on OPC infection induced by a fluconazole resistant clinical isolate of *C. albicans* (fluconazole MIC >128 µg/ml). Again, NCL displayed impressive protection lowering the fungal burden in the oral mucosa by 10 fold versus fluconazole which had no effect on curtailing the infection, as expected (**Fig 8B**). Microscopic analysis of tongue tissue sections stained by Periodic acid-Schiff (PAS) revealed *C. albicans* infection of the superficial epithelial layer similar to pseudomembranous OPC in humans^38^. While placebo gel treated and fluconazole treated mice tongues had widespread infection with abundant tissue penetrating hyphae, NCL treated tongues on the other hand displayed comparatively limited legions, with the infection sites containing a strikingly large numbers of yeasts and pseudohyphae (2X and 10X mag in **Fig 8C**, 40X mag in **S4C**). To validate that NCL-EPO-NP is indeed able to repeal mucosal candidiasis, we additionally tested it independently in a mouse model of vulvovaginal candidiasis. Here too we found that NCL-EPO-NP dose of 20 µg almost completely abrogated mice vaginal infections from the fluconazole resistant *C. albicans* (**S4D**). Fungal burden in NCL-treated mice was at least 1.5 log lower than placebo or fluconazole treated mice. Overall, these *in vivo* studies serve as a testimony to the fact that the anti-hyphae/anti-biofilm activity of NCL-EPO-NP seen *in vitro* is also manifested *in vivo* thereby proving efficacious in prevention of mucosal candidiasis.

**FIGURE 8.**
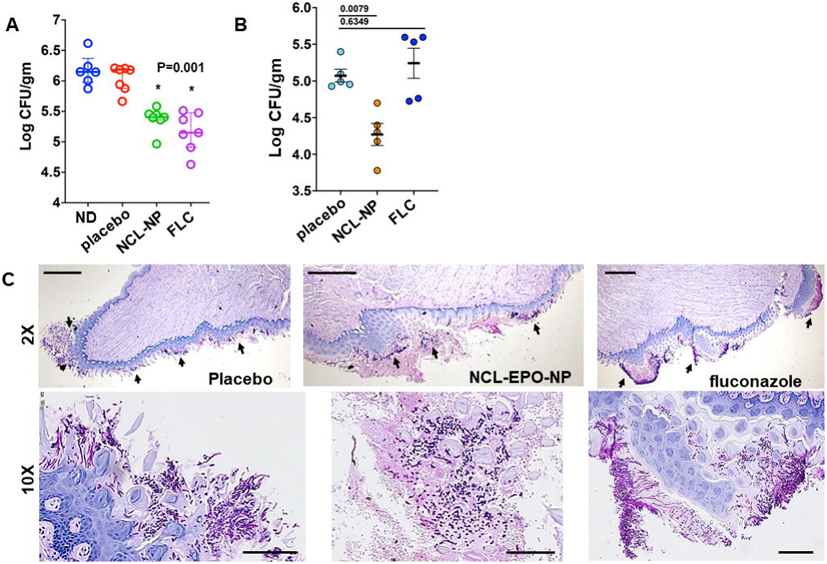
NCL-EPO-NP is efficacious against oropharyngeal candidiasis (OPC). (A) Steroid treated outbred CD1 mice were infected in their oral mucosa with *C. albicans* SC5314, and were treated intra-orally with NCL-EPO-NP gels at 200 µg per dose, twice/day for 4 days. The mice were also treated similarly with gels without NCL as a placebo control, and intraperitoneally with fluconazole on say 3 and 4. Mice sacrificed at day 5 post infection (7 mice per arm) were monitored for fungal CFU in their tongue and presented as log CFU/gm of tissue. (B) Efficacy of NCL-EPO-NP in preventing OPC was also seen during infection by a fluconazole resistant strain of *C. albicans*. (C) The tongues infected with fluconazole resistant strain, and treated with placebo, NCL-EPO-NP or fluconazole were sectioned for histology, stained with PAS, and microscopically visualized at 2X or 10X magnification. Scale bar for upper panel 500 µm, and lower panel 100 µm.

## Discussion

Biofilms display physical gradients of nutrients and gases due to metabolic activity and solute diffusion^40^. The sheer density of cells in an exponentially growing biofilm and the secretion of extrapolymeric substances restrict nutrient diffusion into the innermost layers. This creates a stressful environment especially for the “originator” cells that lay the foundation of biofilm growth on a substrate. Indeed, it is known that *C. albicans* biofilms provide a hypoxic microenvironment, where oxygen concentration decreases steadily from the top to the bottom layers^41^. This happens to such a prominent extent, that deeper layers of *C. albicans* biofilms can sustain growth of obligate anaerobic bacteria. Certainly, the respiratory capacity of *C. albicans* becomes undeniably necessary during this time, with genes associated with mitochondrial respiration upregulated significantly^41^. In fact, the respiratory electron transport chain activity also helps *C. albicans* survive in a glucose-deficient environment, and could prove imperative for adaptation of the fungus to nutrient poor environment in the biofilm core^35^.

We recently published that NDU1, a *C. albicans* mitochondrial protein is required for the assembly and activity of the NADH ubiquinone oxidoreductase Complex I (CI) of the respiratory electron transport chain (ETC)^23^. Absence of NDU1 blocked oxygen uptake, increased ROS accumulation, caused mitochondrial depolarization and prevented *C. albicans* growth on several alternative carbon sources. This phenotype resulted in hypersusceptibility of the fungus to innate immune cells and rendered *Candida* avirulent in a disseminated mouse model. Interestingly, while the NDU1 mutant could develop a robust biofilm, these biofilms were fragile and detached easily from their growth substrate^23^. Although the exact reason of why NDU1 loss causes biofilm detachment is not known, it’s plausible that its absence disallows *C. albicans* to adapt to the hypoxic and nutrient poor environment rampant in the bottom-most biofilm layers. We utilized these phenotypes to identify inhibitors of NDU1 and inducers of early biofilm detachment.

NDU1 mutant grows as adeptly as the WT in media containing glucose, but is completely inviable in acetate as the sole source of carbon. We created a strain of *C. albicans* that overexpressed NDU1 (OE), which can grow both on glucose and acetate. From screening a repurposed library of ~1200 FDA approved inhibitors against this strain, we identified NCL which at a low concentration of 10 µM prevented growth of OE in acetate by 50%, but not in glucose. This indicated that NDU1 could be the target of NCL. It also meant that NCL could be a valuable antifungal molecule considering that the host environment is almost devoid of glucose. Host niches, e.g. bloodstream and tissues only have ~0.05 to 0.1% glucose^42^. FDA approved molecules such as NCL that target respiratory activity of *Candida* which is needed to survive *in vivo*, could be a promising new class of antifungals.

NCL is an anthelmintic indicated in the treatment of tapeworm infections in humans. It is listed on the WHO’s list of essential medicines (file:///C:/Users/puppuluri/Downloads/WHO-MVP-EMP-IAU-2019.06-eng.pdf). It has impressive anti-cancer activity (NCI-lead phase I&II clinical trials against prostrate and colon cancer), and has shown remarkable promise for its broad-spectrum antiviral effect, including on SARS-CoV-2^43–45^. Unfortunately, NCL is minimally absorbed from the gastrointestinal tract—neither the drug nor its metabolites have been recovered from the blood or urine^46^. The poor systemic bioavailability of NCL is because of its limited aqueous solubility of only 0.23 μg/ml. We reckoned that the best use of NCL as an antifungal drug will only be possible if its solubility and absorption characteristics are improved, and the drug can be developed in a way that does not require high doses to enable an antifungal outcome.

One strategy for improving NCM drug solubility has been synthesis of water-soluble analogues^47,48^. However, these new derivatives require comprehensive preclinical/clinical studies to confirm their efficacy and safety and will face several regulatory hurdles prior to approval. An alternate approach is to transform the formulation of this FDA-approved drug by using into clinically viable nanformulations comprised of pharmacologically acceptable excipients to improve its delivery. We exploited a formulation-based approach using an FDA-approved cationic polymethacrylate polymer Eudragit EPO (EPO), for which the pharmacological and toxicological profiles are already known (https://www.fda.gov/media/107564/download). EPO is used in pharmaceutical products as a coating material to achieve taste masking, moisture protection, modified drug release, and for colon targeting ^49–52^. Additionally, EPO, in the form of solid dispersion or nanoparticles (NPs), has been used to improve solubility and oral bioavailability of various hydrophobic drugs including weakly acidic drugs^53–56^.

We and others have previously shown that encapsulation of a weakly acidic and poorly bioavailable anti-inflammatory drug called meloxicam in EPO nanoparticles can improve its efficacy on oral administration with a concomitant reduction in its side effects^31,57^. Given that NCL is weakly acidic and is as hydrophobic as meloxicam, we hypothesized that cationic EPO could encapsulate and stabilize NCL in the NPs via electrostatic interactions. Our preliminary screening involving anionic carboxylic acid terminated polymers such as Eudragit S100, PVAP and cationic EPO confirmed our hypothesis. Polymeric dispersions prepared with Eudragit S100 or PVAP showed rapid precipitation of NCL indicating insufficient interaction and encapsulation of NCL with the anionic polymers. On the contrary, EPO-NCL dispersion showed significant enhancement in color (yellow color) and no signs of NCL precipitation indicating strong interaction and encapsulation of NCL. Our FT-IR and NMR spectroscopic studies confirmed the strong interaction between NCL and EPO at various ratios and corroborated the color enhancement and higher stability observed for NCL-EPO dispersions. While EPO and NCL dispersion showed signs of stability, in order to develop NPs with long-term stability, we evaluated various FDA-approved nonionic surfactants with different chemical structures and HLB values to prepare NCL-EPO-NPs with lowest size and acceptable homogeneity. While nonionic surfactants such as Kolliphor RH40, Vitamin E TPGS and Kolliphor P407 (P407) yielded NCL-EPO-NPs with similar size, we selected P407 as a stabilizer due to its ability to yield higher stability of NCL-EPO-NPs in the initial screening and its capability to enhance penetration and delivery of nanoparticles to the mucosal surfaces ^56,58,59^.

The amount of drug loaded into the NPs significantly impacts the total amount of NPs required for the intended therapeutic effect. Our studies on the loading of NCL into NCL-EPO-NPs revealed a possibility of significantly higher NCL loading into NCL-EPO-NPs (NCL: EPO ratio 4:10 or 40% w/w of EPO) without compromising its size and colloidal stability. Interestingly, hitherto reported NCL nanoformulations involving use of materials such as natural polysaccharides (chitosan and xylan), solid lipids, inorganic materials such as mesoporous silica and hallocyte and biodegradable polymers such as polylactide-co-glycolide (PLGA) to load NCL, were unable to achieve NCL loading > 10% and none of these reports evaluated long-term colloidal stability of NCL nanoformulations ^60–65^. Remarkably, our investigation demonstrated for the first time, high NCL loading in NPs due to strong interaction between NCL and EPO and at least 4-weeks of colloidal stability of NCL-EPO-NP.

The effect of NCL parent drug against *Candida* has previously been studied *in vitro*^66^, against planktonic and biofilm conditions. While that study and ours concur that niclosamide is not fungicidal to *C. albicans* in media containing glucose, we showed that encapsulation in nanoparticles significantly decreases *C. albicans* growth and filamentation in glucose containing media (**Fig 5A,D; S5A**). NCL-EPO-NPs prevented >80% hyphal growth at a striking six to 30 fold lower concentrations than the parent drug alone in RPMI, a hyphal-promoting media (^66^ and **S5B**). These low doses of the drug-loaded nanoparticles could also prevent *C. albicans* invasion of HUVEC cells at a conspicuous 10-15 fold better efficacy than the generic drug alone. Such enhanced anti-invasive activity can be attributed to the complete aqueous solubility, better absorption, and thus improved distribution of NCL due to EPO-nanoparticle encapsulation.

Indeed, prevention of germination and subsequent hyphal formation by NCL-EPO-NPs contributed to its enhanced anti-biofilm activity. Noteworthy was the finding that NCL-EPO-NPs (but not parent NCL; S5C) could disengage developing biofilms early in their development, and penetrate mature biofilms and detach them from their growth surface. Promisingly, NCL-EPO-NPs were not sequestered by the biofilm matrix, a process notoriously known for failure of most antifungal drugs against biofilms^67^. A previous report suggested that interaction between cationic EPO and anionic fluoroquinolones such as ofloxacin could potentiate the permeation and antibacterial activity of ofloxacin by modulation of the electronegative membrane potential of the bacteria leading to alterations in the outer membrane^68^. Given the fact that *C. albicans* exhibits negative membrane potential ^69^, the enhancement of antifungal activity of NCL via encapsulation into EPO NPs could be a combined effect of nanoencapsulation and modulation of the overall negative membrane potential of the *Candida* biofilms.

Bottom-most layers of biofilms are hypoxic and nutrient-starved, subjecting the cells present in the deeper layers to stress, which is combatable by upregulating the fungus’s respiratory machinery. Premature detachment of fully formed biofilms due to NCL-EPO-NP treatment could prospectively be due to loss of viability at the bottommost nutritionally disadvantaged layers of the biofilm, coupled with an early loss of adhesion to substrate due to alterations in the cell wall architecture. We have previously elucidated that loss of NDU1 has a direct impact on *C. albicans* cell wall remodeling and cell membrane integrity (reduction in chitin, cell surface mannan and decrease in ergosterol content) making it susceptible to cell wall and cell membrane perturbing agents^23^. Biofilm disintegration has been considered the gold standard for ridding bacterial biofilm mediated infections^19,70^. Truly, prevention of *C. albicans* biofilms or disintegration of the microbial community early in their development could prove a stellar strategy rendering dissociated cells more susceptible to antifungal drugs^21,70^. The current study is the first to identify a molecule capable of inducing *C. albicans* biofilm detachment.

In an alternative carbon source like acetate, NCL-EPO-NPs was found fungicidal. *C. albicans* can tolerate loss of mitochondrial activity in the presence of glucose, but not in its absence ^35^. Thus, a drug that targets NDU1, an inner mitochondrial membrane protein necessary for electron transport, is expected to be potent under conditions of stress created by the absence of the preferred carbon source. Indeed, NCL-EPO-NPs displayed many of the same phenotypic/biochemical defects as seen in an NDU1 mutant (ROS accumulation, decrease in OCR). Interestingly, unlike *C. albicans*, NCL-EPO-NPs killing activity of *C. auris* was not dependent on carbon source preference. NCL-EPO-NPs prevented planktonic growth of *C. auris* both in glucose and acetate containing media. *C. auris* metabolism favors respiration even in the presence of glucose, as evidenced by enrichment in glycolytic and sugar transporter gene expression during yeast growth and TCA cycle protein enrichment compared to *C. albicans*^71,72^. Respiratory metabolism enhances ATP production and can reduce oxidative stress, also promoting *in vivo* fitness and fluconazole resistance^73^. Thus, NCL-EPO-NP that targets cellular respiration proved lethal to *C. auris* and is poised to be an attractive novel antifungal drug targeting infections caused by this MDR fungus.

The pronounced *in vitro* efficacy of NCL-EPO-NPs against *C. albicans* filamentation, invasion and biofilm growth encouraged us to test this molecule *in vivo*. Previous characterization of NCL pharmacokinetics have revealed poor absorption, which challenges the drug’s efficacy by reducing the dose that reaches the target tissue, precluding its testing in mouse models of bloodstream *Candida* infection ^74^. To overcome this limitation and investigate its antifungal potential *in vivo*, we tested NCL-EPO-NP in two different mucosal models of candidiasis. For uniform delivery of NCL-EPO-NP on intra-oral and intra-vaginal administration, we incorporated NCL-EPO-NP into our previously reported thermosensitive gel composition containing 20% P407 and 1% Poloxamer 188. We and others have demonstrated that thermosensitive P407 hydrogels can be effectively used for the intra-oral and intra-vaginal application of nanoparticles without affecting their inherent properties and release ^36,37,75,76^. Additionally, we have previously shown that nanoparticles stabilized with P407 show significant enhancement in mucosal transport in vivo leading to improved efficacy ^58,59,77^.

Finally, studies have also shown that Poloxamer 407 thermosensitive hydrogel can also interfere with the formation of microbial biofilms ^76,78,79^. Taken together, usage of mucus-penetrating P407 as a stabilizer for EPO-NCL-NPs and P407 thermosensitive gel to deliver NCL-EPO-NPs was deemed advantageous for improving outcome of *in vivo* studies. In both OPC ^38^ as well as the VVC models ^80^, NCL-EPO-NP significantly prevented mucosal fungal burden. This was unmistakably evident by a reduction in oral thrush on the tongue and oral palette of treated mice. The drug prevented *C. albicans* filamentation *in vivo*: histology revealed a remarkable presence of abundant yeast cells in NCL-EPO-NP treated tongue, while the mice treated with placebo or fluconazole displayed a heavy load of filamentous cells. This clearly demonstrated that the efficacy of NCL-EPO-NP *in vitro* against filamentation and biofilm growth could also be deciphered *in vivo* in two biofilm models of mucosal candidiasis.

In summary, we have targeted the respiratory function of *C. albicans* NDU1 to identify an FDA-approved anti-parasitic drug NCL with anti-virulence activity. Additionally, we used an FDA-approved EPO polymer to encapsulate NCL and improve its aqueous solubility, antifungal potency, biofilm penetrability and antifungal activity *in vivo*. This is the first report on the nanoformulation comprised of FDA-approved components that can detach *Candida* biofilms *in vitro* and prevent biofilm infections *in vivo*. Strategies using FDA-approved molecules and materials can undercut the time required for antifungal development and can enhance antimicrobial activity by improving the physiochemical properties of pharmacologically difficult molecules.

## Materials and Methods

### Strains, media, and culture conditions

The following fungal strains were used in this study: *C. albicans* strain SC5314, which is a clinical isolate recovered from a patient with generalized candidiasis ^81^, and two clinical isolates of *Candida* spp. received from the Fungus Testing Laboratory at the University of Texas Health Science Center at San Antonio—fluconazole-resistant *C. albicans* CA6 and CA10. The two *C. auris* isolates CAU-03 and CAU-09 were a kind gift from Shawn Lockhart, Centers for Disease Control and Prevention (CDC). All cultures were maintained by subculture on yeast-peptone-dextrose medium (YPD) at 37°C, and stocks of these cultures were stored in 20% glycerol at −80°C.

### Materials

NCL was purchased from Biosynth-Carbosynth Inc (San Diego, CA). Eudragit^®^ EPO and Eudragit^®^ S100 (Evonik Corporation, Los Angeles, CA), polyvinylacetate phthalate (Colorcon, Inc. West Point, PA), Vitamin E TPGS NF grade (Antares Health Products, Inc., Jonesborough, TN) were received as gift samples. Poloxamer 407 (Kolliphor® P407), Poloxamer 188 (Kolliphor® P188), Kolliphor® ELP, Kolliphor® RH40, Kolliphor® HS15, Kolliphor® PS80, Kolliphor® PS20, were obtained as gift samples from BASF Corporation (Florham Park, NJ). Acetone (AR Grade), scintillation vials (7 ml and 20 ml) and Coumarin C6 were purchased from VWR International (Radnor, PA). Ultra-pure distilled water was used for all experiments.

### HTS

Screening was performed at the Molecular Screening Shared Resource facility at the University of California, Los Angeles. A total of 5 × 10^6^ *C. albicans* yeast cells/ml was suspended in Yeast Nitrogen base containing 2% acetate as an alternative carbon source, and plated (50 µL) into individual 384-well plates using an automated Multidrop 384 system (Thermo Labsystems). The New Prestwick Chemical Library consisting of 1,233 drugs was used to pin one compound per well at 10 µM final concentration, using a Biomek FX liquid handler. Twenty four hours later the plates were scanned with a Flex Station II 384-well plate reader (Molecular Devices) to measure turbidity (OD600) of the wells. Molecules displaying >50% reduction in turbidity compared to control non-drug-treated wells (MIC50) were considered primary “hits.”

### Preliminary studies to assess stabilization potential of Eudragit EPO

A preliminary study was carried out to evaluate the potential of Eudragit EPO (cationic polymethacrylate), polyvinylacetate phthalate (PVAP; anionic polycarboxylate), and Eudragit S100 (anionic polymethacrylate) to stabilize weakly acidic NCL. Briefly, NCL (10 mg) and Eudragit EPO/PVAP/Eudragit S100 (100 mg) were transferred to a 20 ml scintillation vial and the mixture was dissolved in 10 ml of acetone to obtain a uniform solution. Ultra-pure water (10 ml) was transferred to a 50 ml clean and dry beaker and the beaker was placed on a magnetic stirrer. Organic solution containing NCL, and the polymer was added dropwise to the aqueous solution and the mixture was stirred at 600 rpm in a fume hood for 3 hours to evaporate acetone. The resultant dispersion was observed for signs of precipitation over a period of time.

### Evaluation of concentration-dependent molecular interaction between EPO and NCL using Fourier-transform infrared (FT-IR) spectroscopy and nuclear magnetic resonance (NMR) spectroscopy

For the spectroscopic studies, NCL and EPO in different weight ratios (% w/w NCL: EPO = 1:10, 2:10, 3:10, and 4:10) were weighed and transferred to a scintillation vial. The mixture was dissolved in acetone to obtain a uniform solution. Acetone was evaporated using a rotary evaporator (Scilogex, LLC, Rocky Hill, CT) to obtain a solid dispersion of NCL and EPO. The solid dispersion was further dried overnight using a vacuum oven (Fisher Scientific, Waltham, MA). The NCL-EPO solid dispersions, EPO, and NCL were evaluated using the FTIR spectrometer (Nicolet iS10, Thermo Scientific, Waltham, MA) equipped with a diamond attenuated total reflection unit. The FTIR spectra were obtained in the transmission mode from 4000 to 500 cm^−1^. The 1H NMR spectra of NCL-EPO solid dispersions, EPO, and NCL were obtained using a Bruker Avance Digital 400 MHz NMR spectrometer (Bruker, Billerica, MA) coupled to a BACS 1 automatic sample changer. The spectrometer is equipped with a 5-mm PABBO BB-1H/D Z-GRD probe. The spectra of the samples were recorded in (400 μl) deuterated chloroform (CDCl_3_, Acros Organics, 99.5% D, Waltham, MA) with an average of 16 scans. The chemical shifts were reported in ppm using residual deuterated solvent peaks as an internal reference for 1H NMR: CDCl_3_ (7.26 ppm).

### Development and characterization of NCL loaded Eudragit EPO nanoparticles (NCL-EPO-NPs): Screening of surfactants for their ability to form NCL-EPO-NPs

Various FDA-approved surfactants such as Poloxamer 407 (P407), Kolliphor® ELP (ELP), Kolliphor® RH40 (RH40), Kolliphor® HS15 (HS15), Kolliphor® PS80 (PS80), Kolliphor® PS20 (PS20), and Vitamin E TPGS (TPGS) were screened for the development of NCL-EPO-NPs. Briefly, Eudragit EPO (50 mg), NCL (5 mg), and surfactant (200 mg) were transferred to a 20 mL scintillation vial and the contents were completely dissolved in 10 ml acetone by vortexing (organic phase). The aqueous phase consisting of 10 mL ultra-pure water was transferred to a clean 50 mL beaker. The beaker containing water was placed on a multi-point magnetic stirrer (IKA Works) and the stirrer was set at 600 rpm. The organic phase was slowly added to the aqueous phase to avoid any splashing and to allow the formation of NPs. The stirring was continued for at least 3 hours in a fume hood to allow for complete evaporation of the organic solvent. The nanoparticles synthesis experiment was carried out in triplicate. The size and polydispersity index of various NCL-EPO-NPs were evaluated using Litesizer 500 particle analyzer (Anton-Paar USA, Inc., Torrance, CA). The surfactant that yielded NCL-EPO-NPs with the lowest size and polydispersity index was selected for further experiments.

### Preparation of NCL-EPO-NPs with different NCL loading and their characterization

Previously carried out EPO-NCL molecular interaction studies formed the basis for the preparation of NCL-EPO-NPs with different NCL loading. Briefly, NCL (10, 20, 30, or 40 mg), EPO (100 mg), and P407 (400 mg) were transferred to a 20 ml scintillation vial and dissolved in 10 ml of acetone to obtain a homogenous organic phase. The aqueous phase consisting of 10 mL ultra-pure water was transferred to a clean 50 mL beaker. The remaining procedure for the preparation of NCL-EPO-NPs was similar to that described in the earlier section. The size, polydispersity index, and zeta potential of various NCL-EPO-NPs were evaluated using Litesizer 500 particle analyzer (Anton-Paar USA, Inc., Torrance, CA). The nanoparticle synthesis experiment was carried out in triplicate. For TEM imaging, NCL-EPO-NPs with NCL loading of 40% w/w EPO were selected. Briefly, samples were applied to glow-discharged 200 mesh carbon-coated Formvar-coated copper grids and allowed to dry. Grids were viewed on a Hitachi HT7700 TEM at 100kV and photographed with an AMT XR-41B 2k x 2k CCD camera.

### Determination of the encapsulation efficiency of NCL in NCL-EPO-NPs

NCL-EPO-NPs dispersion (0.5 ml) was transferred to the Amicon Ultra-0.5 device (3 kDa membrane; Fisher Scientific, Waltham, MA). The Amicon-Ultra centrifugal filter containing NCL-EPO-NPs was centrifuged for 15 min at 14000 rpm (Thermo Scientific, Legend Micro 17R centrifuge). The filtrate was then diluted 10-fold with methanol and NCL concentration in the filtrate was measured at 340 nm using a UV-Vis spectrophotometer. The % encapsulation efficiency (EE) was calculated by the following equation:

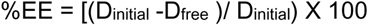

where ‘D_initial_’ is the amount of NCL/mL of EPO-NCL-NPs dispersion and D_free_ is the amount of NCL/mL of the filtrate obtained by the centrifugation of NCL-EPO-NPs.

### UV-Vis spectrophotometric determination of NCL from NCL-EPO-NPs

Briefly, NCL (10 mg) and EPO (25 mg) were transferred to a 100 ml volumetric flask and the contents were dissolved in 100 ml methanol to obtain a stock solution of NCL (NCL concentration 100 µg/ml). The stock solution was suitably diluted with methanol:water (99:1) to obtain various NCL concentrations (2, 2.5, 5, 7.5, 10, 15, and 20 µg/ml). The absorbance of solutions containing various NCL concentration was measured at 340 nm using Agilent Carey 60 UV-Vis Spectrophotometer. NCL calibration curve was obtained (n=3) by plotting a graph of NCL absorbance Vs NCL concentration. For the determination of NCL content from NCL-EPO-NPs, the NPs were diluted 100-1000-fold with methanol to obtain NCL concentration with the range established by NCL calibration curve and the absorbance of the NCL was measured at 340 nm.

### Evaluation of molecular interaction between NCL and EPO in NCL-EPO-NPs using fluorescence spectroscopy, FT-IR, and 1H-NMR

Fluorescence spectroscopy was used to determine the molecular interaction between EPO and NCL in NCL-EPO-NPs dispersion. Briefly, NCL-EPO-NPs with NCL loading of 40% w/w EPO were diluted with water to obtain a solution with NCL concentration of 25 µg/ml. NCL hydroalcoholic solution (NCL concentration: 25 µg/ml) was used as a control. The fluorescence spectra of NCL from NCL-EPO-NPs dispersion and control NCL solution was evaluated using Shimadzu RF 5301 Spectrofluorometer using an excitation wavelength of 310 nm for NCL solution and 330 nm for NCL-EPO-NPs. For the NCL solution, the excitation wavelength of 310 nm was used to minimize Raman scattering associated with the solvent which interfered with the fluorescence analysis. For FT-IR and NMR characterization, NCL-EPO-NPs were freeze dried. Briefly, NCL-EPO-NPs containing NCL loading of 40% w/w were transferred to a scintillation vial and the contents in the vial were flash frozen using liquid nitrogen. The frozen vial was transferred to a freeze drying flask and the freeze drying flask containing NCL-EPO-NPs samples were loaded on the Labconco FreeZone 12 Plus freeze dryer preset at −80°C temperature and <0.12 mb pressure. After 24 h of lyophilization, the samples were removed from the freeze dryer and the vial containing freeze dried NCL-EPO-NPs were stored in a vacuum desiccator until further use. The freeze dried NCL-EPO-NPs were evaluated using FT-IR and NMR to ascertain molecular interaction between NCL and EPO in the nanoparticles.

### Evaluation of long-term physical and chemical stability of EPO-NCL-NPs

The optimized EPO-NCL-NPs containing different loading of NCL were evaluated for the 4-week long physical stability (size, polydispersity index, zeta potential and pH) whereas NCL-EPO-NPs containing 40% w/w NCL was evaluated for 4-weeks long chemical stability using fluorescence spectrophotometer.

### Preparation of Coumarin-6 encapsulated polymeric NPs

Briefly, Eudragit EPO (100 mg) and Kolliphor P407 (400 mg) were transferred to a 20 mL scintillation vial along with 9.95 ml of acetone. The contents in the scintillation vial were vigorously vortexed to obtain a clear solution (organic phase). Coumarin-6 (Acros, Organics) was dissolved in acetone at a concentration of 2 mg/mL and added to the organic phase containing polymers to obtain a final concentration of 0.01 mg/mL. The polymeric NPs were prepared as described earlier. The unencapsulated Coumarin-6 in the NPs was removed using Amicon Ultra-0.5 Centrifugal Filter Devices (Amicon Ultra 10K device). Briefly, nanoparticle dispersion (0.5 mL) was transferred to the Amicon-Ultra-0.5 device and centrifuged for 15 min at 14000 rpm (Thermo Scientific, Legend Micro 17R centrifuge). The NPs were washed twice with ultra-pure distilled water to get rid of free Coumarin-6. The filtrate was discarded and the concentrated Coumarin-6 encapsulated polymeric NPs (~100 µL) were collected in a separate tube by inverting and centrifuging the Amicon Ultra filter for 2 min at 1000 rpm. The concentrated NPs were then diluted to their original volume by adding ~400 μL of distilled water, the tube was covered with aluminum foil and stored until further use.

### Development of NCL loaded Eudragit EPO nanoparticles (NCL-EPO-NPs) thermosensitive gel

Briefly, 10 mL of blank EPO-NPs were taken in a 20 mL scintillation vial. Kolliphor P407 (20% w/v) and Kolliphor 188 (1% w/v) were added into it and vortexed. The vial was kept in the refrigerator (4° C) for 3 h to ensure total dissolution of surfactants. The same procedure was performed to obtain NCL-EPO-NPs thermosensitive gel of the optimized batch.

### Dose-response assays

Dose-response assay of NCL-EPO-NP against planktonically grown fungi was performed in agreement with the CLSI M27-A3 (for yeast) reference standards for antifungal susceptibility testing^82^. Each drug was used in the concentration range of 0.6 µg/ml to 64 µg/ml, and the MIC of NCL-EPO-NP in media containing 2% acetate was compared to media containing 2% glucose. Inhibition of planktonic growth or filamentation due to drug treatment was also visualized and imaged using bright-field microscopy.

### Biofilm growth and drug susceptibility testing

Biofilms of *Candida* spp. were developed in 96-well microtiter plates, and susceptibility of the biofilm cells to NCL-EPO-NP was determined as described previously ^83^. Biofilms were initiated in either the presence or absence of the drugs, or the drugs were tested on 48-h preformed biofilms. Inhibition of biofilm growth was measured by a standard colorimetric assay (XTT) that measures metabolic activity of the biofilm cells ^83^. Absorbance at 490 nm was measured using an automated plate reader. Biofilm disruption by the NCL-EPO-NP were also visualized by bright field microscopy. *C. albicans* 48 h mature biofilms were also treated with 2 µg/ml coumarin-6 encapsulated NCL-EPO-NP for 24 h. Biofilms were additionally stained with 25 µg/ml concanavalin A to visualize the cells in red. Confocal microscopy with z-stacking feature was performed and images were collected at two wavelengths, 488 nm (GFP) and 594 nm (ConA) as published previously^84^.

### ROS measurement

Intracellular ROS production was detected by staining cells with 5 μM MitoSOX Red (Life Technologies, Frederick, Maryland, USA) in DMSO. Cells from 25-ml cultures grown for 6 h at 30°C in the presence and absence of NCL-EPO-NP in YPD medium were collected and washed twice with PBS. The pellets were suspended to 1 × 10^6^ cells in 1 ml of PBS and treated with or without MitoSOX Red for 45 min at 30°C in the dark. Cell fluorescence in the presence of DMSO alone was used to verify that background fluorescence was similar per strain. Cells from each MitoSOX-treated sample were collected and washed twice with PBS after staining, and mean fluorescence for ROS was quantified.

### Oxygen consumption rate assay

OCR were measured under a Seahorse instrument (Seahorse Bioscience, Massachusetts, USA) according to the manufacturer’s instructions. Cells were plated in poly-d-lysine coated XF96 microplates at a density of 5×10^4^ cells/well in 100 µL volume of unsupplemented DMEM (no glucose) with or without NCL-EPO-NP. Samples were run with three technical replicates per condition. The plates were centrifuged at 1000 rpm for 4 minutes without use of the centrifuge break. After centrifugation plate was incubated for 5 minutes before being loaded into a Seahorse XF96 Extracellular Flux Analyzer (Agilent). Basal respiration was measured over six hours with 36 ten-minute protocol cycles including a: 3 minute mix, 4 minute wait, and 3 minute measurement. After six hours, the Complex I and Complex III electron transport chain inhibitors rotenone and antimycin A were injected from Ports A and B at a final concentration of 5µM to determine non-mitochondrial respiration. Data analysis included subtraction of non-mitochondrial respiration from basal respiration measurements before averaging the last 4 basal measurements.

### Mammalian cell damage assays

Primary human umbilical vascular endothelial cells (HUVECs) were used to determine the cytotoxicity and efficacy of the NCL-EPO-NP in preventing damage by *C. albcians*. HUVECs were isolated and propagated by the method of Jaffe *et al.*^85^. The cells were grown in M-199 (Gibco, Grand Island, NY) supplemented with 10% fetal bovine serum, 10% defined bovine calf serum, and 2 nM l-glutamine, with penicillin and streptomycin. Second- or third-passage endothelial cells were grown on collagen matrix on 96-well microtiter plates. Treatment with NCL-EPO-NP was conducted in M-199 medium. Different concentrations of the nanoparticles were introduced into the cell lines and the wells were also added with 1×10^5^ *C. albicans* yeast cells. Plates were incubated for 6 h at 37°C in 5% CO_2_. Damage to HUVEC cells were measured using the chromium release assay well published previously by our group^25,86^.

### Animal models

All animal related study procedures were compliant with the Animal Welfare Act, the Guide for the Care and Use of Laboratory Animals, and the Office of Laboratory Animal Welfare and were conducted under an IACUC approved protocol 31443-03 by The Lundquist Institute at Harbor-UCLA Medical Center.

### Mouse model of oropharyngeal candidiasis (OPC)

Male, 6-week-old CD1 mice were purchased from Taconic. OPC was induced in mice as described previously^38^. Mice were injected subcutaneously with cortisone acetate (225 mg/kg of body weight) on days −1, 1, and 3 relative to infection. For inoculation, the animals were sedated, and a swab saturated with 2 × 10^7^ *C. albicans* cells was placed sublingually for 75 min. Mice were either treated intraperitoneally with 10 mg/kg fluconazole on days 3 and 4 after infection. Some mice were treated intra-orally with 200 µg of NCL-EPO-NP on days 1-4. Mice were sacrificed on day 5 after infection. The tongues were harvested, weighed, homogenized for 30 sec, and quantitatively cultured. Some tongues were fixed in zinc-buffered formalin, embedded in paraffin, sectioned, and stained with periodic acid-Schiff (PAS).

### Mouse model of vulvovaginal candidiasis

Mice were infected vaginally using a previously well-published protocol^80^. Five to 6 week old female CD1 mice were injected with 0.1 mL of β-Estradiol 17-valerate in sesame oil, subcutaneously in the back of the neck on day −3 and +3 relative to infection using a needle size of 20G - 27G. For inoculation, the mice were anaesthetized by intraperitoneal (i.p) injection of a mixture of ketamine (82.5 mg/kg) and xylazine (6 mg/kg). Next, 20 μL of PBS containing 5 × 10^6^ *C. albicans* blastospores was injected into the vaginal lumen. A group of mice were also treated intra-vaginally with fluconazole or NCL-EPO-NP (fluconazole: 10 mg/kg intraperitoneally once on days 3 and 4 after infection; NCL-EPO-NP: 200 µg intra-vaginally on days 1-4). On day 5 post infection, vaginas and ~1 cm of each uterine horn was dissected, homogenized, and quantitatively cultured.

## Data availability

The raw data used to reproduce these findings will be shared on request during the review period.

## Supporting information

Supplemental only

## Acknowledgements

We would like to thank the following agencies for their financial support to carry out this project: NIH NIAID R01AI141794 awarded to PU and NIH NEI R24EY033598 awarded to AD. The authors would also like to thank BASF Corp, USA, Evonik Industries and Colorcon, Inc. for providing excipients for our research work.

**SUPPLEMENTAL 1:** Results of the HTS and shortlisting of hits

**SUPPLEMENTAL 2:** UV-Visible spectra of Niclosamide (NCL-10 ppm), NCL-EPO NPs (equivalent to 10 ppm NCL) & EPO blank NPs (A). Calibration curve for NCL using UV-Vis Spectroscopy (n=3 ± S.D.) (B). Summary of the characteristic FT-IR peaks representing the key functional groups in Eudragit EPO, Niclosamide (NCL), Kolliphor P407 and NCL-EPO-NPs (freeze dried) (C). ^1^H NMR chemical shifts observed in freeze dried NCL-EPO-NPs in comparison with EPO and NCL (D). Evaluation of particle size, polydispersity index (PDI) and zeta potential of NCL-EPO-NPs containing different loading of NCL (% w/w EPO) on long-term storage at room temperature. (Data expressed as mean ± S.D.; n =3) (E).

**SUPPLEMENTAL 3:** NCL-EPO-NPs prevent planktonic *C. albicans* growth in acetate (A), protect HUVEC from damage (B), penetrate *C. albicans* biofilms (C), and increase ROS activity in the fungus (D)

**SUPPLEMENTAL 4:** NCL-EPO-NPs reduce oxygen consumption rate in *C. albicans* a time course analysis (A); NCL-EPO-NPs can be formulated as gel (B); Histology pictures from the OPC model, 40X magnification, scale=20 µm (C); Fungal burden in vagina after drug treatment (D)

**SUPPLEMENTAL 5:** NCL-EPO-NP reduces *C. albicans* growth and filamentation at ~0.5 µg/ml in both YP+1% glucose or RPMI medium {when cells remain in the yeast form they fall to the bottom of the round-bottom wells forming a ring that is easily identifiable by the naked eye; filamentation instead displays a fuzzy circumference^87^}. NCL parent is ineffective even at >32 µg/ml (A,B). Parent NCL or nanoparticle control (NP) did not detach biofilms while NCL-EPO-NP detached preformed biofilms at concentrations as low as 0.5-1 µg/ml (C). NCL-EPO-NP abrogates filamentation at 1 µg/ml, while parent NCL does not, and filaments as efficiently as the no drug control.

