## Supplementary figures and images for "Niclosamide loaded Eudragit EPO nanoparticles show enhanced *Candida* biofilm penetration, trigger biofilm detachment and protect from mucosal candidiasis"

### S2.TIF

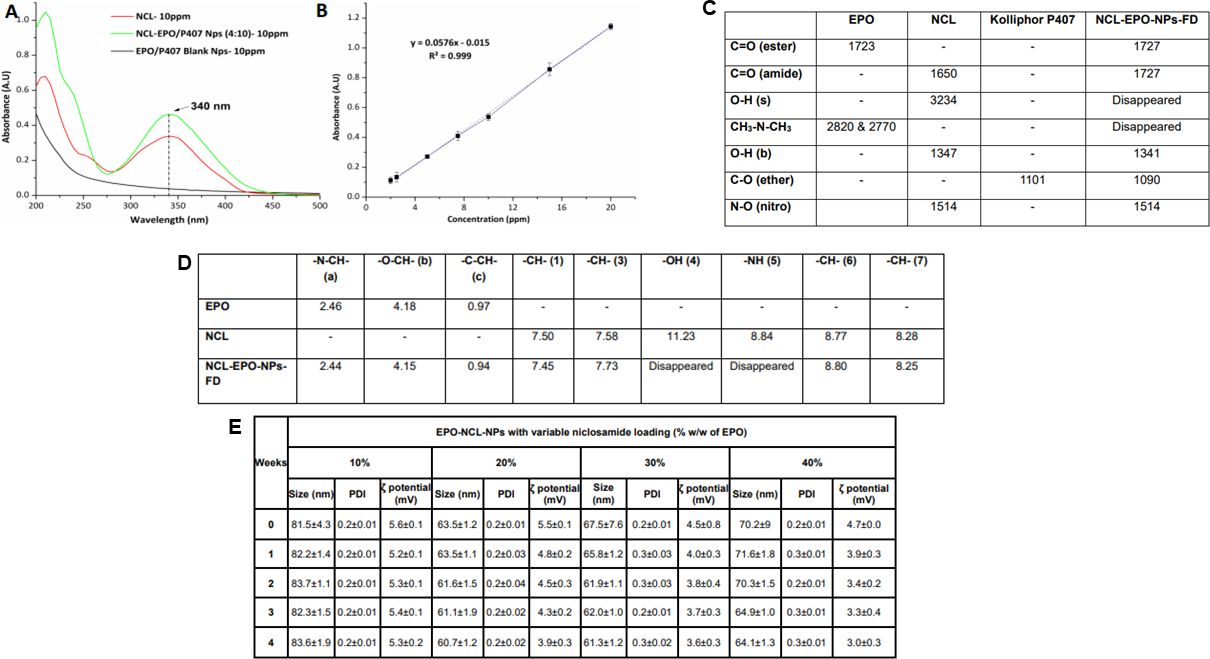

### S3.TIF

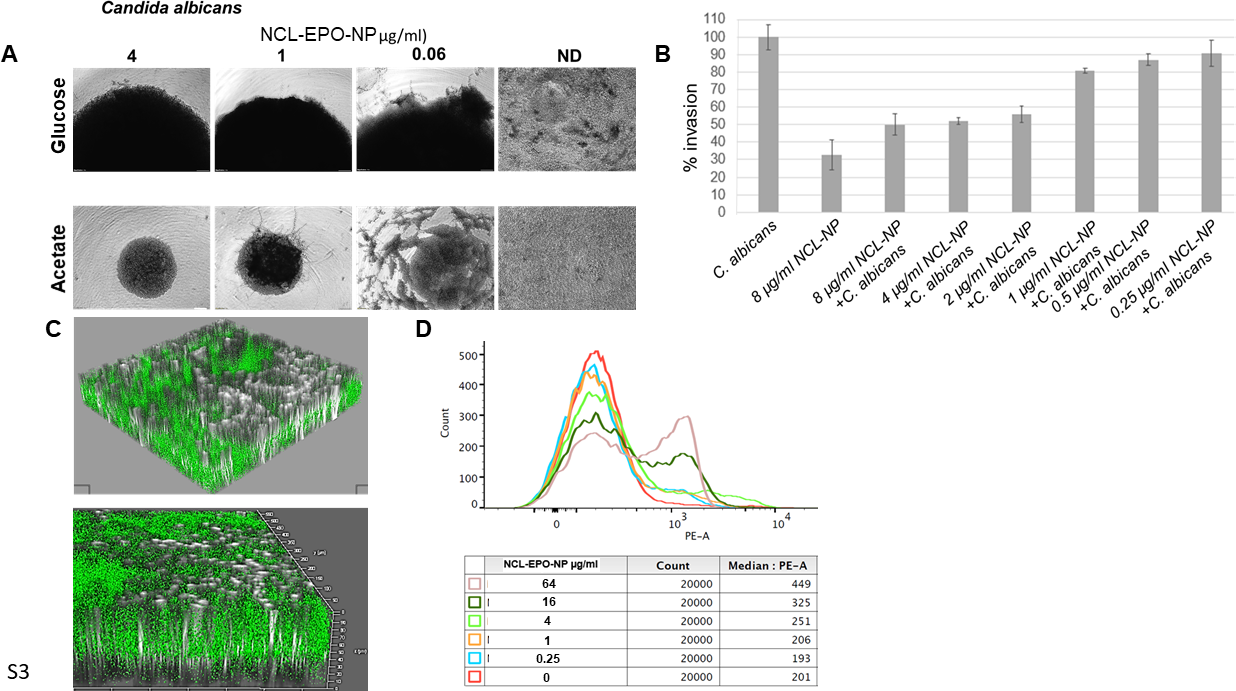

### S4.TIF

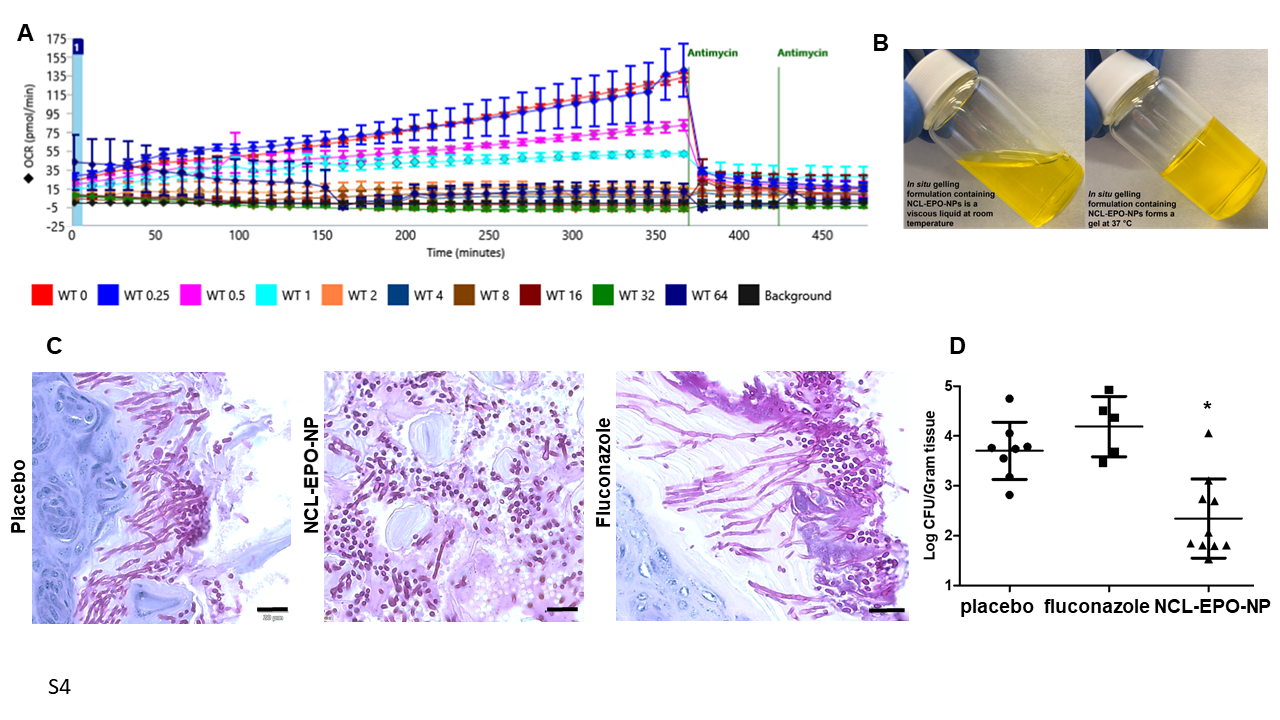

### S5.TIF

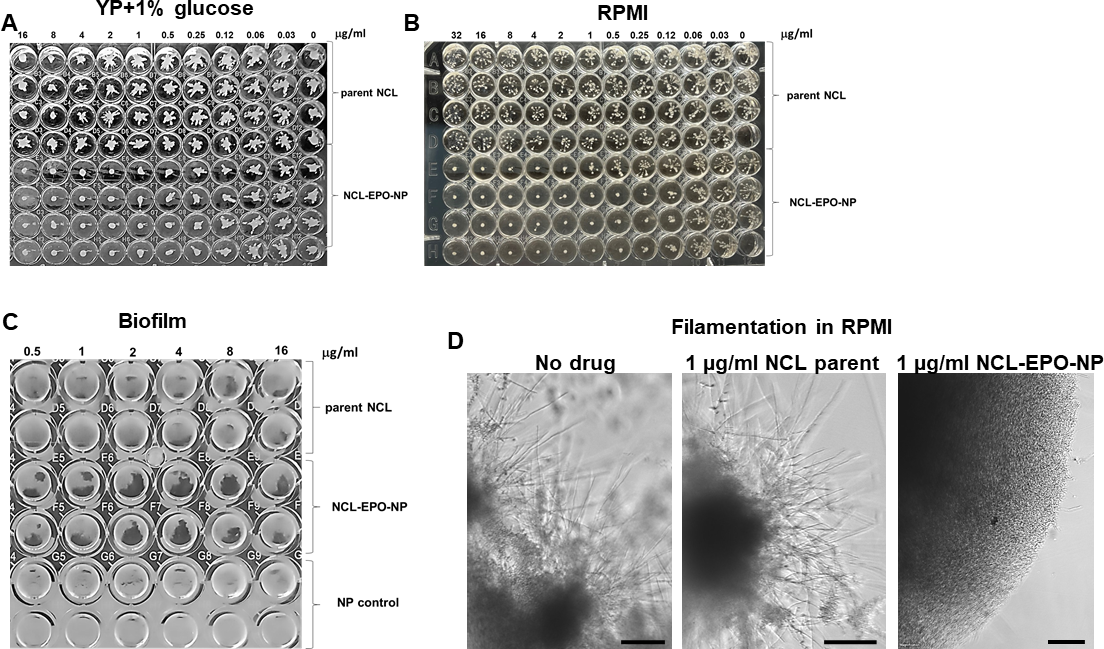
